# Does plant –microbe’s common Inter Simple Sequence Repeat lead to a new recombination? A novel mechanism of pathogenicity

**DOI:** 10.1101/2021.01.21.427556

**Authors:** Purvi M. Rakhashiya, Pritesh P. Bhatt, Vrinda S. Thaker

**Affiliations:** Department of Biosciences, Saurashtra University, Rajkot 360 005, Gujarat, and Vimal Research Society for Agrobiotech, Vekohouse, 80ft Road, Agi area Rajkot, Gujarat, India

**Keywords:** DNA hybridization, ISSR markers, Mango (*Mangifera indica* L.), Mechanism of pathogenecity, Pathogens, Recombination

## Abstract

**Abstract:** A total of eight varieties of the mango from an orchard were studied using molecular markers to understand the host-pathogen interaction. From the infected leaves of the plant, a total of the 8 bacterial pathogens (*Exiguobacterium arabatum, Pseudomonas mendocina, Pantoea dispersa, Bacillus sp. Pantoea ananatis, Micrococcous luteus, Microbacterium_sp., Enterobacter cloacae*) were isolated and identified. All the host varieties of mango were distinguished for the genetic diversity using the Inter Simple Sequence Repeat (ISSR) DNA markers. This set of ISSR marker primers were also used for the mango pathogens. PCR amplification of the ISSR primers showed polymorphic and monomorphic band patterns in the host plants and in their pathogens. The monomorphic band generated by PCR amplification in the host and in the pathogen, by the common primer, is selected and used for PCR hybridization technique. PCR products obtained from the host, pathogen and hybridization were cloned, sequenced and compared. A multiple sequence alignment of these sequences revealed that the product of hybridization PCR was mixture of host and pathogen sequences. On this basis, we hypothesize a possibility for the recombination of host-microbes DNA as one of the mechanisms of pathogenicity for the plant pathogens using hybrid PCR technique. The possible mechanism of recombination for plant host and its pathogen is discussed.

**Highlights:** Inter Simple Sequence Repeat markers used to (i) Fingerprint the pathogens and their host (mango) and (ii) for study of the possibilities for the recombination as mechanism of pathogenicity.

## Introduction

Mechanisms of pathogenicity of phytopathogens are being extensively studied in Gram-negative and some Gram-positive bacteria (Baker *et al.*, 2010; Venturi and Fuqua, 2013). These studies revealed that pathogens contain suits of genes including various transporter systems (Chen *et al.*, 2010; Rakhashiya *et al.*, 2016; Wilkens, 2015), secretion systems (TSS), virulence and associated metabolism (Abby *et al.*, 2016; Chang *et al.*, 2014; Green and Mecsas, 2016). In pathogens, from the cytoplasm through the inner membrane or the outer membrane to the extracellular space, the secretion of proteins requires dedicated types of machinery, i.e. type I to V secretion pathways (Thanassi and Hultgren, 2000). The distinction in these pathways from each other is mainly by the presence or absence of a signal peptide on the secreted protein and by the mode of translocation steps. Using these types of machinery, pathogen gets an entry into the host cell and colonizes there. We report in this study, a probable new mechanism of pathogenicity of the bacterial pathogen *(M. luteus)* in its plant host (*Mangifera indica* L. Varieties Totapuri). We hypothesized that common repeat region of the genome of host and pathogen may also offer an opportunity to new recombination of genomic DNAs and hence the DNA marker technique with repeat region amplification is selected for this experiment.

Many fruit species are screened for the desired characters with the molecular markers such as banana (Brown *et al.*, 2009), litchi (Chundet *et al.*, 2007), pummel (Uzun *et al.*, 2013), grape (Vafaee *et al.*, 2017) and sweet orange (Dehesdtani *et al.*, 2007). Inter-simple sequence repeats (ISSRs) is a PCR-based DNA marker technique shows reproducible results and it is widely used for the identification of plant and animal species but the references in bacteriology are rather scanty. It is found in the genome flanked by microsatellite sequences (Simple Sequence Repeat) regions. Random DNA segments in the genome can be PCR-amplified by using arbitrarily designed primers. Such experiments usually produce multiple DNA fragments (each of which is considered a locus) in a single reaction. Thus without the need to first know the DNA sequences of the target regions, allowing the generation of a large number of loci across the genome of any species. The technique is mainly useful for the understanding evaluation of genetic diversity, cultivar identification and phylogenetic relationship of the studied organisms. The simplicity of ISSR markers predetermines them for gene tagging and it is also used with QTL analysis (Agarwal *et al.*, 2008; Khaled and Hamam, 2015; Malik *et al.*, 2014). ISSR with its ease of application, we envision that it can be explored to comprehend the host-microbe interaction.

Mango (*Mangifera indica* L.), ‘the king of the fruits’ is an economically important plant with high nutritional value (Sivakumar *et al.*, 2011; Tharanathan *et al.*, 2006). Nine different host varieties of mango from the orchard are studied for the ISSR primers. The same primers are used for the mango pathogens and hypothesized a possibility for the recombination of host-microbes DNA and one of the mechanisms of pathogenicity for the plant pathogens. To the best of our knowledge, this the first study to DNA fingerprint plant and its pathogenic microorganism using ISSR markers. This DNA profiling technology is used in this experiment to (i) enlarge the number of molecular markers that are suitable in the molecular characterization of either host (mango varieties) or their isolated pathogens and (ii) examination of the genetic relationship between host and pathogens together (if any) which can be judged by common oligonucleotide patterns and hence the possibilities of new recombination.

## Materials and Methods

### Collection of plant material and isolation of the organisms

Healthy and infected leaves (leaf with black spot) of different varieties of *Mangifera indica* L. were collected from the Mango orchard near Rajkot city (22.30°N, 70.78°E). The pathogenic Gram’s positive and negative bacteria used in this study were isolated from the infected leaves of their host plants (Table 1). The infected leaves of mango were washed continuously under tap water for 3 h then it was treated with 0.1% HgCl_2_ for 15 min. Further, it was washed thrice in sterile distilled water (Rakhashiya *et al.*, 2015a; Rakhashiya *et al.*, 2015b). The Infected part of the leaf was inoculated on N-agar plates and plates were incubated at 37 °C for 24-48 h.

**Table 1.** Host *Mangifera indica* L. Varieties (1-9) and their pathogens (M1-M8)

| No. | Plant Variety | No. | Bacteria | Accession no. |
| --- | --- | --- | --- | --- |
| 1 | Ratnagiri Happus | M1 | *Exiguobacterium arabatum* | KP836352 |
| 2 | Jamadar | M2 | *Pseudomonas mendocina* | KP836353 |
| 3 | Nylon | M3 | *Pantoea dispersa* | KP836354 |
| 4 | Nil Franso | M4 | *Bacillus sp.* | - |
| 5 | Kesar | M5 | *Pantoea ananatis* | JF802201 |
| 6 | Totapuri | M6 | *Micrococcous luteus* | KM454981 |
| 7 | Manglori badam | M7 | *Microbacterium_sp.* | KM365269 |
| 8 | Betisha Tirupati | M8 | *Enterobacter cloacae* | KP8366103 |
| 9 | New variety – The variety used for Koch's' postulate Test |  |  |  |

### Identification of bacterial strains

The pathogen strains were identified by morphological analysis (Gram’s staining), biochemical tests (according to Bergey’s Manual of Systematic Bacteriology), and molecular (16S rDNA) technique. For molecular techniques, DNA isolation, 16S rDNA amplification, and 16S rDNA sequencing were carried out in-house for the identification. Genomic DNA of mango pathogens was isolated according to (Chudasama and Thaker, 2014) and the plant DNA of healthy leaves was isolated using the protocol of (Mandaliya *et al.*, 2010). 16S rRNA gene of each pathogen was amplified in PCR (Viriti™ Thermal Cycler, Applied Biosystems) using universal primer sets (8F (5′-AGAGTTTGATCCTGGCTCAG -3’) and 1525R (5’-ACGGCTACCTTGTTACGACTT -3’). Genomic DNA used as a template for amplification of 16S rRNA gene (35 cycles, denaturation − 94 °C for 5 minutes, annealing − 47.5 °C (and 52 °C for 8F, 1525R) for 1 minute, final extension − 72 °C for 12 minutes). 16S rDNA gene sequencing was performed on 3130 Genetic Analyzer (Applied Biosystems, U.S.A.). These sequences were used for molecular identification with NCBI BLAST tool. The nucleotide sequences of studied pathogens are submitted to NCBI Genbank Database using BankIt submission tool (Table 1).

#### ISSR primer amplification

DNA of pathogens and/or host plant varieties was subjected to ISSR primer amplification. A set of 30 different ISSR primers were used for this study (Table 2). The total volume of 25 μl contained 2.5 μl 10x buffer (10mM Tris-HCl pH 9.0, 50 mM KCl, 0.1% Triton X-100), 2.5 μl 25 mM MgCl2, 1μl 10mM deoxynucleotide, 10 μM primer and 1 μl of Taq DNA polymerase, 2 μl DNA template. PCR reaction was performed using the TM Thermal cycler with 40 cycles of denaturation at 92 °C for 2 min., annealing (36-60 °C, Table 2) for 1 min., extension was done at 72 °C for 7 min. Amplified DNA fragments were analyzed through 2% agarose gel electrophoresis. All PCR reactions are repeated at least twice.

**Table 2.** ISSR primers used in this study for host and pathogens.

| Sr. no | Primer name | sequence | Temperature (°c) |
| --- | --- | --- | --- |
| *I-1* | UBC803 | (AT)8C | 36 |
| *I-2* | UBC808 | (AG)8C | 48 |
| *I-3* | UBC814 | (CT)8A | 48 |
| *I-4* | UBC817 | (CA)8A | 44.2 |
| *I-5* | UBC822 | (TC)8A | 48 |
| *I-6* | UBC826 | (AC)8C | 48 |
| *I-7* | UBC834 | (AG)8TT | 44.2 |
| *I-8* | UBC840 | (GA)8TT | 44.2 |
| *I-9* | UBC862 | (AGC)6 | 48 |
| *I-10* | UBC864 | (ATG)6 | 48 |
| *I-11* | UBC867 | (GGC)6 | 48 |
| *I-12* | UBC868 | (GAA)6 | 48 |
| *I-13* | UBC872 | (GATA)4 | 48 |
| *I-14* | UBC873 | (GACA)6 | 48 |
| *I-15* | UBC876 | (GATA)2(GACA)2 | 48 |
| *I-16* | UBC880 | (GGAGA)3 | 48 |
| *I-17* | UBC881 | (GGGGT)3G | 48 |
| *I-18* | UBC884 | HBH(AG)7 | 48 |
| *I-19* | UBC885 | BHB(GA)7 | 48 |
| *I-20* | UBC886 | VDV(CT)7 | 48 |
| *I-21* | UBC888 | BDB(CA)7 | 48 |
| *I-22* | UBC889 | DBD(AC)7 | 48 |
| *I-23* | UBC891 | HVH(TG)7 | 48 |
| *I-24* | ISSR1 | (AGC)5GC | 55 |
| *I-25* | ISSR2 | (CA)7AC | 44.2 |
| *I-26* | ISSR3 | (GT)7AC | 44.2 |
| *I-27* | ISSR4 | GCA(GA)7 | 47.2 |
| *I-28* | ISSR5 | (GA)9C | 51.7 |
| *I-29* | ISSR6 | (GA)9A | 49.5 |
| *I-30* | ISSR7 | (CG)8C | 60 |

#### PCR hybridization

This method involves PCR amplification using ISSR primer- UBC 814 (This primer generates monomorphic band in host and pathogen). Initially, PCR amplification was performed in pathogen and host plant individually (First PCR). Amplified PCR product of these PCRs were mixed and used as DNA template for new PCR (Second PCR) amplification. The products obtained after first and second PCR amplifications were observed on 2% agarose gel electrophoresis.

#### Cloning

The amplified products of first PCR (pathogen and host) and second PCR (hybrid) were cloned and sequenced. UBC 814 primer generated monomorphic bands were cloned by using ‘InsTAclone PCR Cloning Kit’ as suggested by the manufacturer (Thermo Scientific, USA). As per the protocol, direct one-step cloning using TA system for PCR products with 3’-dA overhangs to dT overhangs of vector was done. *E. coli* strain DH5α with lacZΔM15 mutation was used for the transformation of cloned vectors. The plates were incubated for 16 hours at 37°C temperature and screened for blue/white colonies to check successful transformation. The white colonies were selected for the plasmid isolation by alkaline lysis method (Sambrook *et al.*, 1989).

#### Plasmids insert amplification and Nucleotide sequencing

The desired cloned fragment from plasmid was amplified using M13 primers. The inserted products (of pathogen/ host/ hybrid) were cycle sequenced using Applied Biosystems BigDye Terminator v3.1 Cycle Sequencing Chemistry on ABI Genetic Analyzer 3130 (Fernandez-Carvajal *et al.*, 2009). Sequences from the ISSR monomorphic bands of host and pathogen and hybrid assay were compared. The multiple sequence alignment (MSA) was performed on CLC Main Workbench7 to study the variation patterns in the studied sequences.

#### Data analysis

PCR products obtained using ISSR primers were scored for the band pattern analysis according to (Vafaee *et al.*, 2017) for each primer and % polymorphism and % monomorphic was calculated. Phylogenetic analyses was performed using unweighted pair-group method with arithmetic averages (UPGMA), as suggested by (Sneath and Sokal, 1973) and dendrogram was constructed using Free Tree and CLC workbench 7.0.2. Software.

## Results and Discussion

In the present study, we observed eight different mango varieties infected by different pathogenic bacteria exhibited almost similar morphological symptoms but 16s RNA sequences results showed that the pathogens are different (Table 1). The common symptoms observed in all varieties is leaf black The 9^th^ variety was taken as a healthy plant and all pathogens were individually tested and confirmed (Koch’s postulate) on this variety (data is not presented). These hosts and pathogens were evaluated for the set of the ISSR primers to distinguish among hosts and/or pathogens and to understand the mechanism of recombination (if any) between their chromosomal DNA.

### ISSR studies in mango

For the first approach, total 30 ISSR primers were used to assess the genetic variability among 8 pathogens and 9 varieties of mango. From these, 12 primers amplified in all plant varieties and their pathogens. The amplification patterns obtained by the 12 primers showed typical ISSR banding pattern (Fig.1). The set of primers used throughout this study allowed more abundant and reproducible banding profiles. Amplification of ISSR markers yielded 701 fragments in the plant. The highest number of bands observed with I-20 (UBC 886). The studies on genetic diversity of the plants using the ISSR primer technique is well documented (Mandaliya *et al.*, 2015; Mansour *et al.*, 2010; Monpara *et al.*, 2017) including some cultivars of mango (Gajera *et al.*, 2011; Samal *et al.*, 2012). In the present study, all host varieties containing narrow genetic variations can be distinguished with the studied ISSR primers.

**Fig.1.**
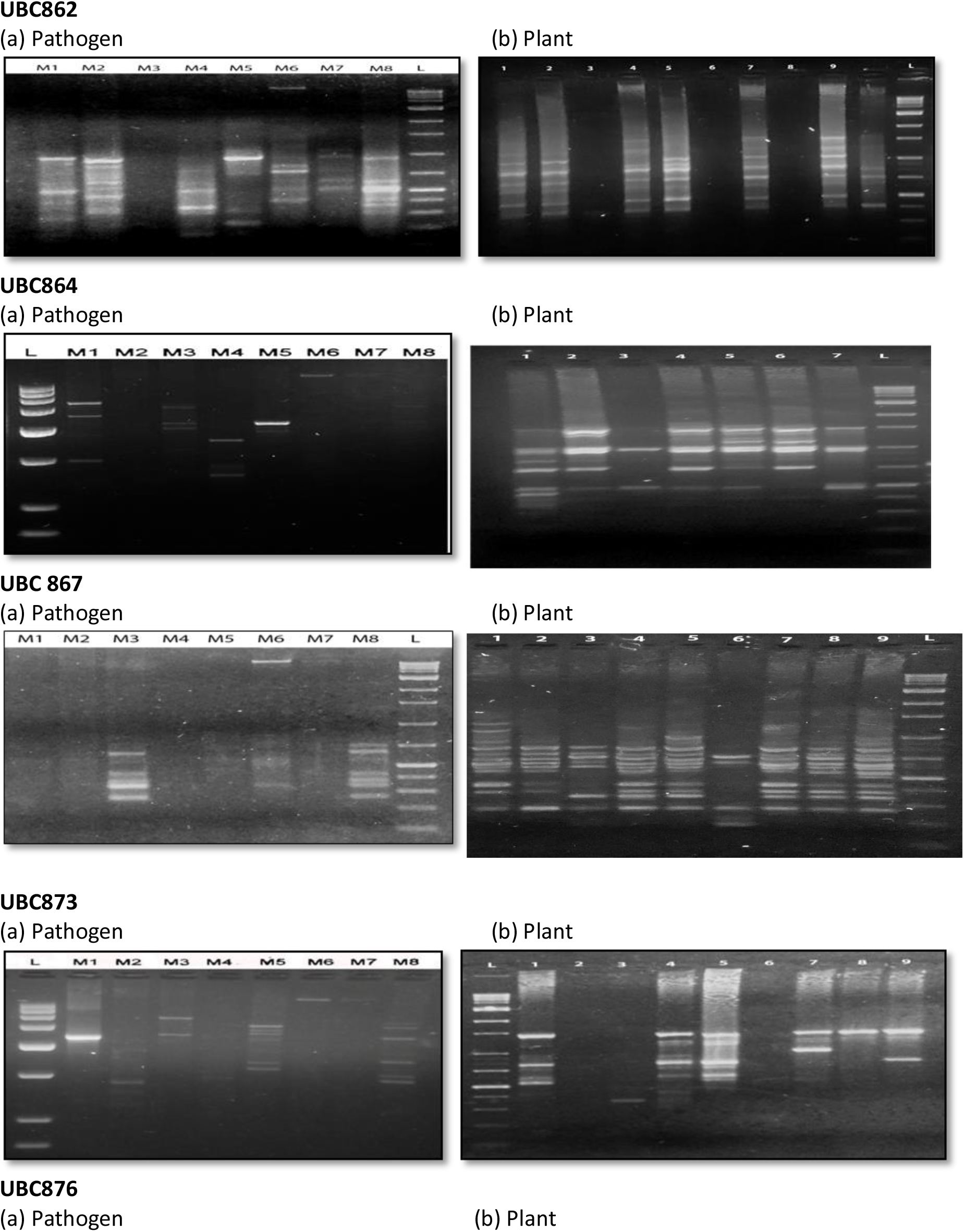
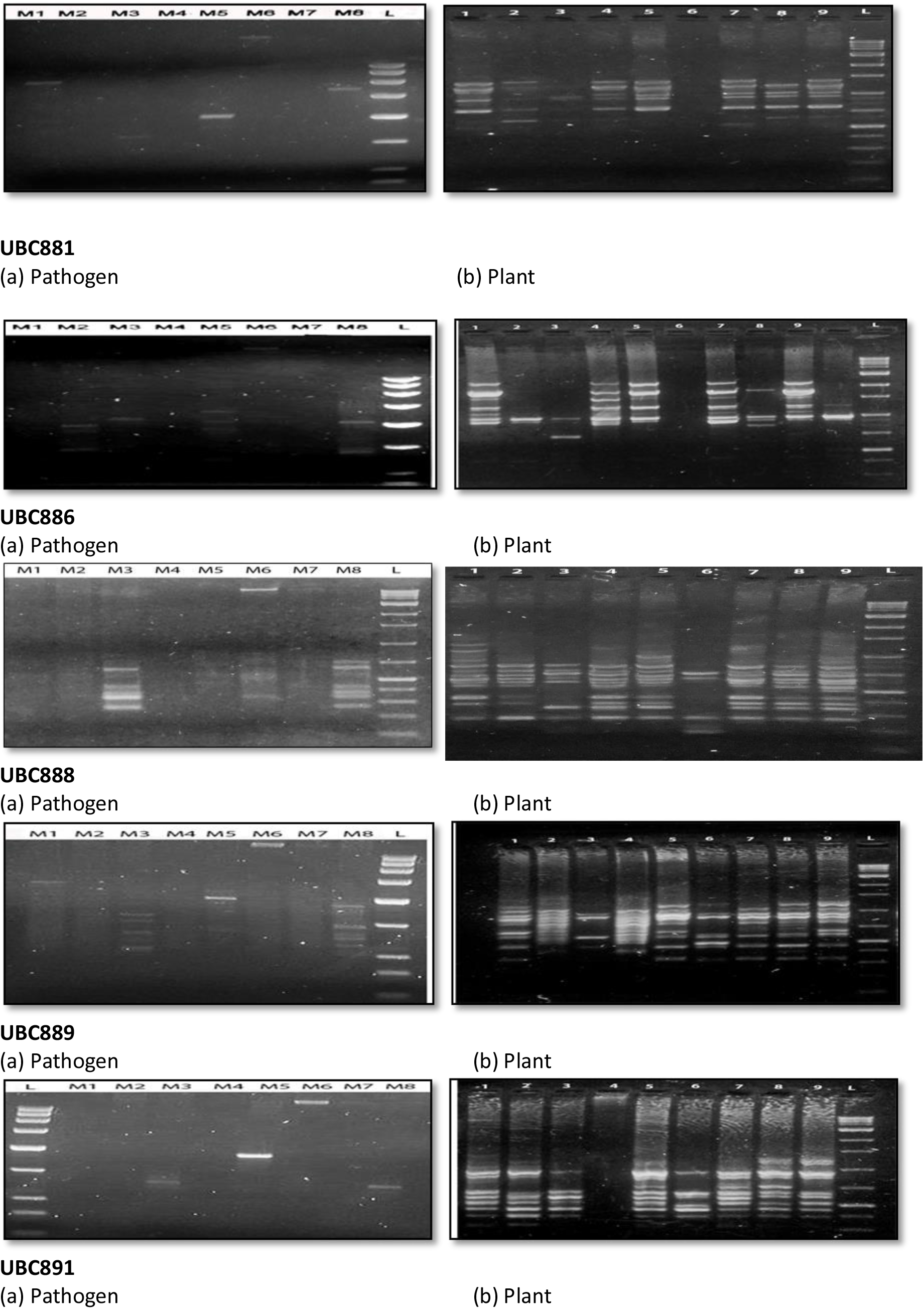
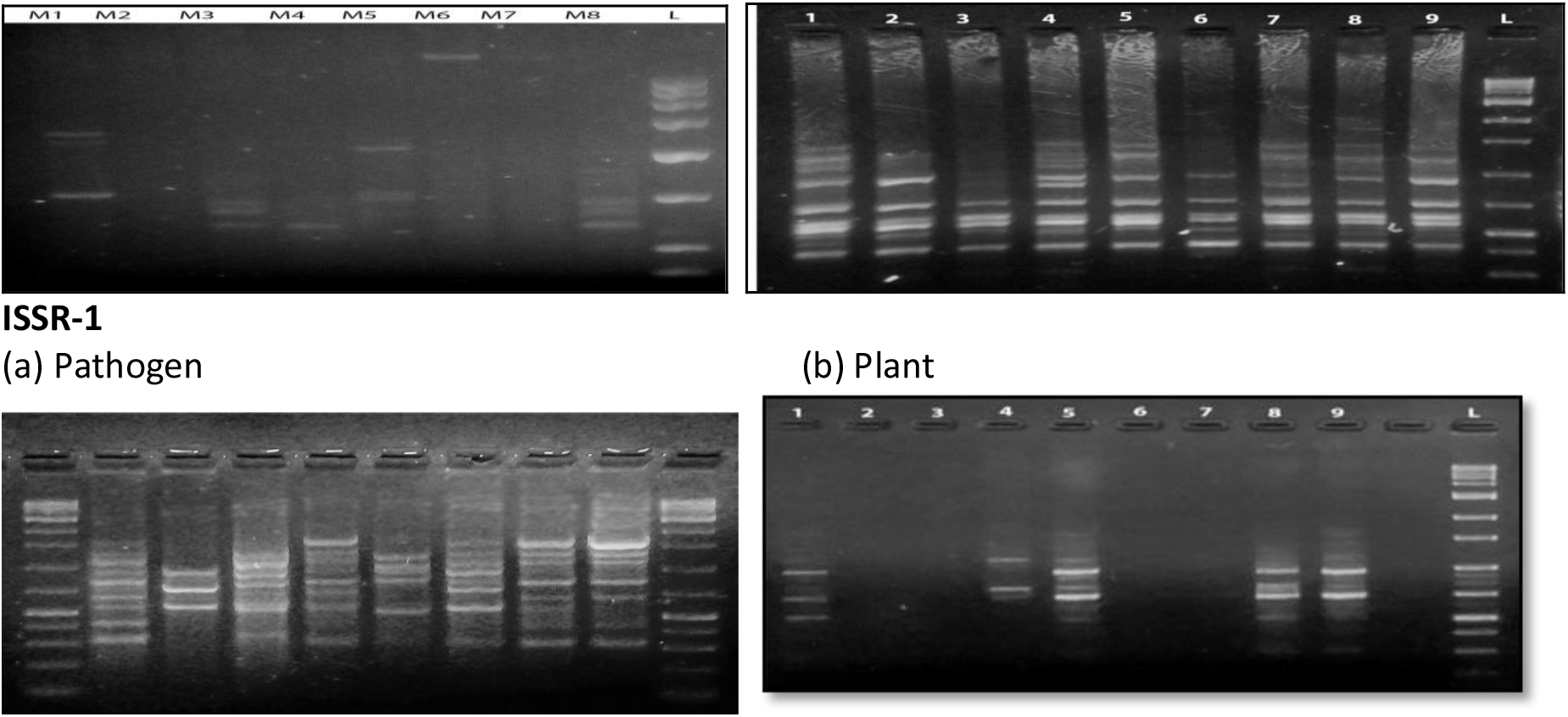
ISSR band patterns in pathogens (M1-M8) and host plants (1-9). Below each ISSR primer code, (a) pathogen and (b) plant DNA amplification profile.

### ISSR primer amplification in pathogens

DNA segments were also amplified in the pathogens revealed unique observations (Fig. 1). Notably, this technique in bacteria demonstrated that all the pathogens produced different fragments amplified and provided uniqueness to the results. ISSR-PCR generated reproducible banding patterns, regardless of repeated PCR assays, the concentration of template DNA or the subculture. Total 194 fragments are generated in bacteria, in this, the highest number of polymorphic bands obtained by ISSR primer I-24 (ISSR 1). Interestingly, all the organisms showed at least one common ISSR primer amplification with the 9^th^ variety (Fig. 1). Each ISSR bands were scored as present (1) or absent (0) and as an independent character regardless of its intensity. The bacterial dendrogram clearly distinguished eight bacterial pathogens into 3 major clades; but band pattern showed clear distinction among all the pathogens (Fig. 1) these results support to distinguish the bacterial diversity using the technique. ISSR marker studies for bacteria are scanty in the literature; on plant and its pathogens, this is the probably first report. In the present study, ISSR profiling helps to distinguish and fingerprint the pathogens of the same host plant when the disease symptoms are common.

### PCR hybrid extension

In this paper, we have selected a pathogen, *M. luteus* (M6), and its host variety, (*M. indica* L. Totapuri), for the hybridization assay as both generate monomorphic band with ISSR primer UBC814 (Fig. 3). The hybrid extension by PCR as used in the present study involves three separate PCRs; the two DNA fragments produced in the ISSR reactions in bacteria and plant, and the mixture to form the hybrid template for the next stage. It was assumed that hybrid template will have some primer binding sites common to plant and bacteria and the middle portion of the sequences may/ may not be similar and subsequent PCR should generate recombinant DNA sequences. ISSR marker (UBC 814) gave monomorphic band patterns in the results (Fig.2); which can be cloned and used as a molecular marker for the individual pathogen/plant. In addition, the primer generates monomorphic band in host and pathogen is tested for the hybrid PCR and sequencing. For this, the monomorphic bands were taken and denatured at 95°C in PCR and allowed to cool down at room temperature. In fact, the polymorphic bands generated more opportunities for recombination of two different DNAs; but here selected monomorphic band, considering the ease of the assay to understand the mechanism. The PCR mixture used as template DNA with the same primer set was amplified and sequenced (Fig. 4). Further, the hybrid PCR with the DNA templates of pathogen and host generated chimeric DNA which might have joined in the PCR cycles within short stretches of identity. The characteristic of the end products is an important concern in understanding recombination reactions; for example, are their overall structures simply or multiply exchanged? Further, is the joining homologous or illegitimate? The newly formed sequence is accurate or mutagenic? In this deference, the most important feature of the newly generated sequence is that either template is applied for the point of new recombination (Fig. 3, 4). These results inferences our understanding of the mechanism(s) of genetic recombination in the host and pathogen DNAs. Such interactions between similar (homologous) DNA sequences play an important biological role in host-pathogen interactions.

**Fig.2.**
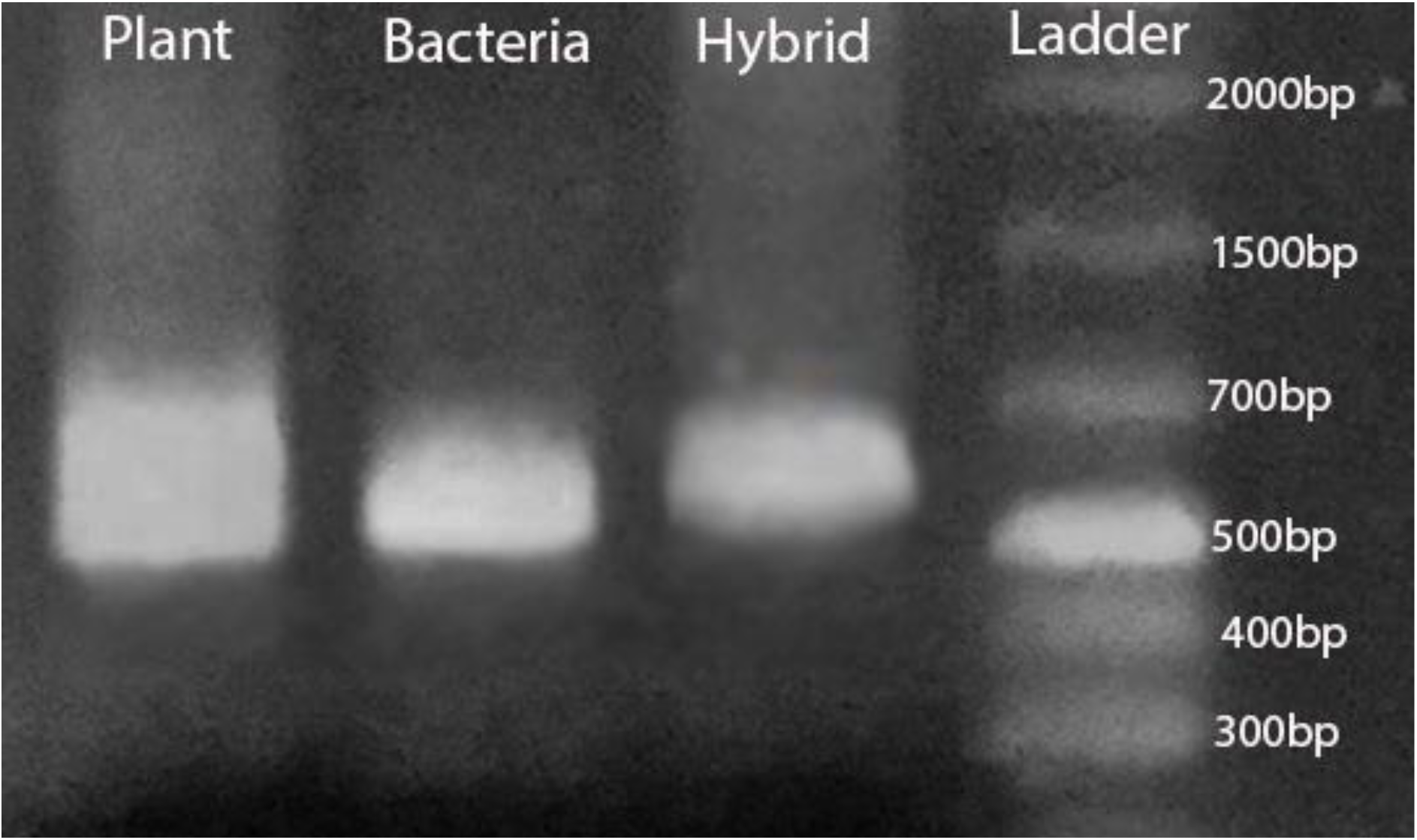
Monomorphic band pattern in Host (plant), Pathogen (bacteria) and mixed DNA template (hybrid) using USB814 primer.

**Fig.3.**
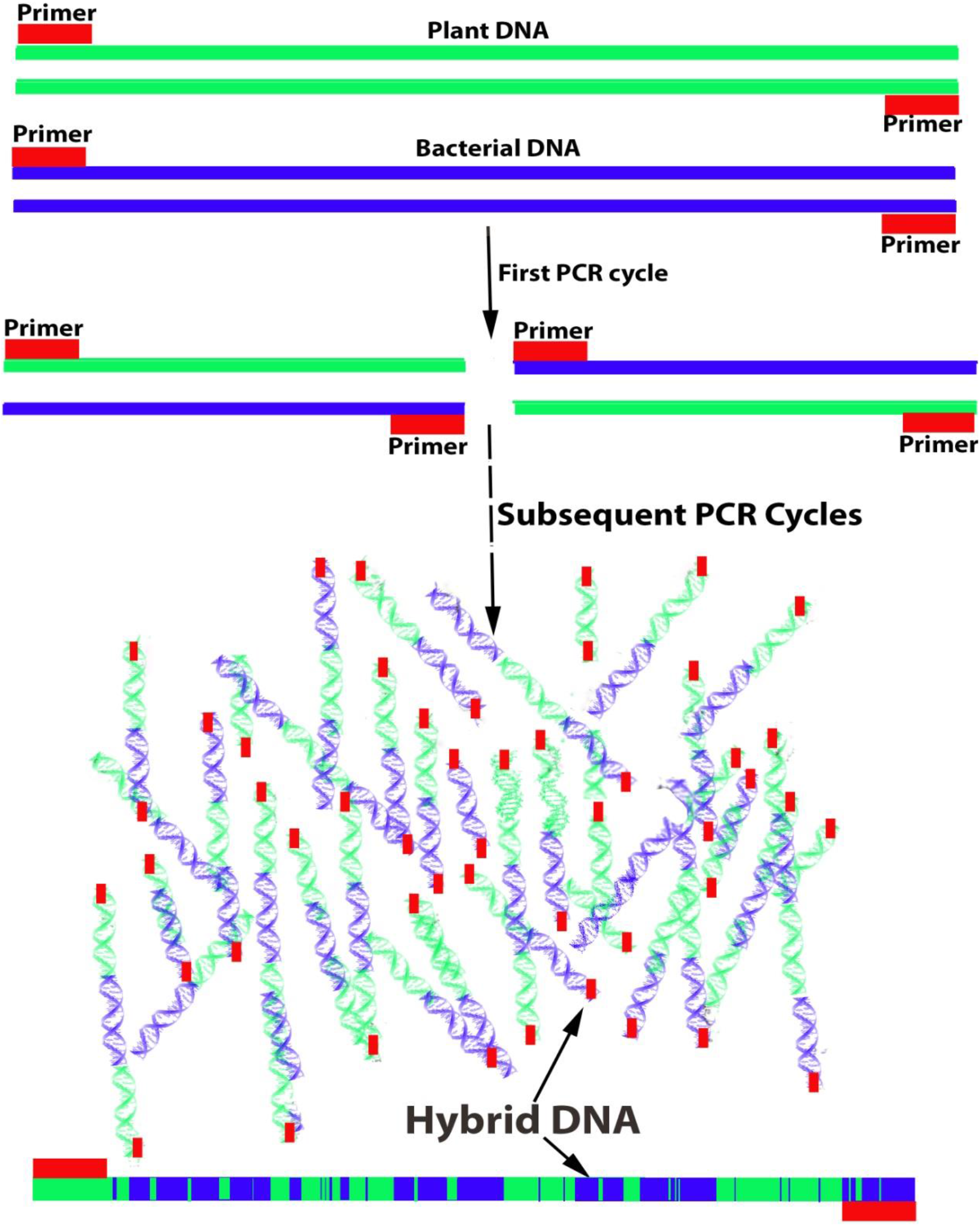
A schematic diagram of recombination of host and pathogen DNA.

**Fig.4.**
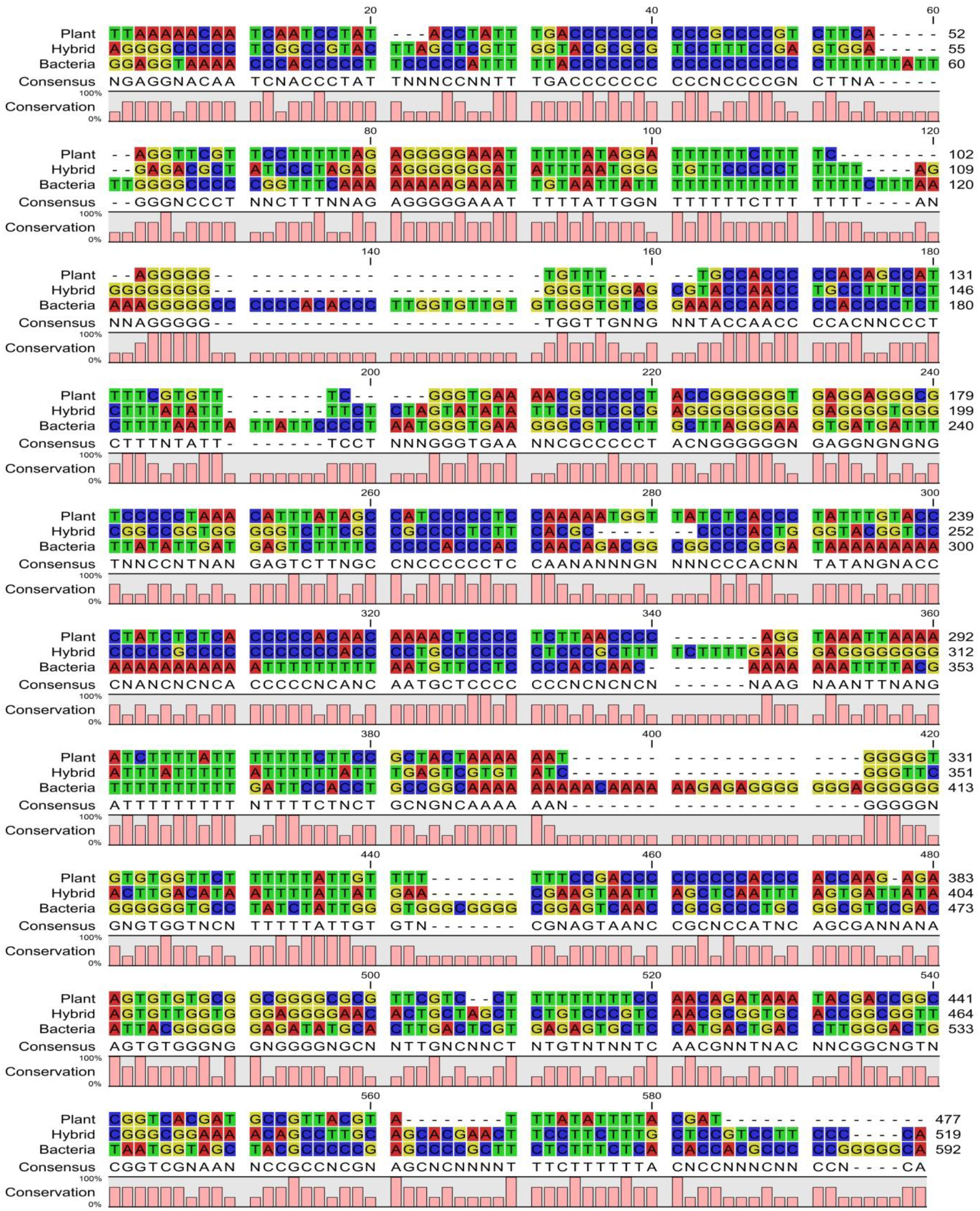
Sequence alignment of host plant, hybrid (host and pathogen DNA templates) and bacteria (pathogen) after PCR amplification and sequencing using ISSR primer USB814.

It revealed that sequence of plant and pathogen are different in alignment results; but the hybrid sequence showed a mixture of both, randomly. We proposed that the splicing overlap and mutations in the hybrid sequence observed attributed to the ISSR technique that uses single primer for forward and reverse amplification. We also observed that the length of the monomorphic band of host and pathogen were not exactly equal and that each PCR cycle increased the probabilities of variations (Fig.3, 4). Such these results suggest that the possibilities of recombination in host-pathogen cannot be ruled out.

## Acknowledgements

Authors are thankful to UGC-CAS Department of Biosciences Saurashtra University, Department of higher education, and State Government of Gujarat for the financial support.

## Author contributions

Rakhashiya PM and Bhatt PP have done experiment and data analysis, Thaker VS given the experiment design the script writing.

